# Preclinical Safety Evaluation of Vernolac, a Commercially Available Polyherbal Nutraceutical Comprising *Vernonia zeylanica, Nigella sativa, Hemidesmus indica, Smilax glabra, and Leucas zeylanica,* in Wistar Rats

**DOI:** 10.64898/2025.12.27.696713

**Authors:** Shalini K. Wijerathne, Epitawala Arachchige Oshadi, Sandani De Vass Gunawardana, Ashein Kothalawala, Tharindu Lakmal Dissanayake, Gayani Ranaweera, Arumugam Murugananthan, Prasanna Galhena, Kanishka Senathilake, Sameera R. Samarakoon

## Abstract

Vernolac is a commercially available polyherbal formulation comprising *Vernonia zeylanica* aerial parts, *Nigella sativa s*eeds, *Hemidesmus indica* roots, *Smilax glabra* rhizome, and *Leucas zeylanica* aerial parts. Although previous *in vitro* studies have demonstrated anticancer potential of Vernolac and its active ingredients, safety data are available only for some of the plant ingredients of Vernolac. In the present study, acute and 28-day repeat-dose toxicity of Vernolac were evaluated in Wistar rats following OECD guidelines 420 and 407, respectively. Acute toxic effect was investigated during 14 days after administering a single oral dose of 2000 mg/kg to 10 animals (5 males, 5 female) which was followed by repeat-dose study where a human equivalent therapeutic dose HED (165 mg/kg/day), a mid-dose (2x HED, 330mg/kg/day) and a high dose (4x HED, 660 mg/kg/day) were administered separately to a group of 10 fresh animals (5 male, 5 female) for 28 days. In both acute and repeat-dose studies, no morbidity, mortality, changes in food and water intake, relative organ weights, microscopic changes in organs, hematological changes, or clinical signs of toxicity were observed. In the acute study, significant differences appeared only in AST and ALT levels in males, indicating the liver may be a target organ of toxicity at extremely high doses. In 28-days repeated dose study, a significant reduction in ALT was observed only in females receiving high doses, with no changes in males. In summary, a dose up to four times the therapeutic daily dose is non-toxic in Wistar rats over a 28-day period.

## 1. Introduction

Cancer continues to be a major global health challenge and remains among the leading causes of mortality worldwide. Despite substantial advances in early diagnostic techniques and therapeutic interventions, the overall burden of cancer is steadily increasing [1,2]. Conventional treatment modalities, including surgery, chemotherapy, radiotherapy, and targeted therapies, constitute the cornerstone of cancer management; however, their clinical utility is often limited by significant adverse effects [3]. In addition, the development of therapeutic resistance poses a major obstacle, frequently resulting in disease progression and treatment failure [4]. Collectively, these limitations highlight the critical need for innovative cancer therapies that are both efficacious and better tolerated.

Natural products have attracted considerable interest in the field of cancer therapy owing to their wide-ranging therapeutic potential [5]. A diverse array of bioactive compounds sourced from plants, marine organisms, fungi, and microorganisms has demonstrated anticancer activity through the regulation of critical cellular pathways, including apoptosis, cell proliferation, angiogenesis, and immune modulation [6]. Within this framework, nutraceuticals have gained prominence as potential complementary agents to standard anticancer treatments [7]. Their appeal lies in their ability to act on multiple molecular targets while minimizing damage to normal tissues, thereby supporting their suitability for prolonged application in cancer management strategies [8].

Polyherbal nutraceutical formulations are of particular interest due to their multi-component and multi-target nature, which may enhance therapeutic efficacy through synergistic mechanisms [9]. However, this same complexity raises critical safety concerns, as interactions among bioactive phytochemicals can alter pharmacological and toxicological profiles in ways that cannot be predicted from studies of individual plant constituents [10]. Therefore, regulatory authorities and international guidelines emphasize the necessity of rigorous preclinical safety evaluation of herbal formulations, especially those intended for repeated or prolonged human consumption [10].

Vernolac is a polyherbal nutraceutical capsule manufactured by Fadna Life Sciences (Pvt) Ltd (106/6B, Araliya Uyana, Depanama, Pannipitiya, Sri Lanka), currently available in the Sri Lankan market for the management of cancer. The formulation comprises five medicinal plants with ethnomedical relevance, namely *Vernonia zeylanica* (aerial parts), *Nigella sativa* (seeds), *Smilax glabra* (rhizome), *Leucas zeylanica* (aerial parts), and *Hemidesmus indicus* (roots) [11]. The formulation is currently used alongside conventional cancer therapies, with claims related to supporting anticancer activity, reducing oxidative stress, and improving patient well-being during long-term treatment [12].

A polyherbal decoction consisting of *N. sativa* seeds, *H. indicus* roots, and *S. glabra* rhizomes in equal proportions has been traditionally prescribed for the management of cancer by a lineage of Ayurvedic physicians in Sri Lanka, underscoring its long-standing ethnopharmacological significance in oncology [13]. Experimental studies have demonstrated that this decoction confers marked protection against chemically induced hepatocarcinogenesis in rat models, with no evidence of significant toxicity [14,15]. Additionally, the formulation has demonstrated notable cytotoxic activity against human hepatoma (HepG2) cells *in vitro* [13,16]. Mechanistic investigations suggest that its anticancer effects are mediated through multiple pathways, including attenuation of oxidative stress [17], suppression of inflammatory responses [18], and modulation of apoptotic signaling via the regulation of pro- and anti-apoptotic gene expression [13]. A recent study demonstrated that a supercritical CO₂ extract of Vernolac exhibits potent and selective anticancer activity against cancer stem cell–like NTERA-2 cl.D1 cells, inducing apoptosis, oxidative stress, and suppression of migration while sparing non-cancerous cells, thereby supporting its potential as a targeted anticancer nutraceutical [12]. An integrative network pharmacology and *in vitro* study revealed that Vernolac contains multiple bioactive phytochemicals targeting key cancer-related pathways, with predicted actions involving apoptosis induction, immune modulation, and antiproliferative mechanisms [19].

Despite this growing body of efficacy-oriented evidence, a critical knowledge gap remains: the safety profile of Vernolac as a combined polyherbal formulation has not been systematically evaluated *in vivo*. Importantly, cytotoxic and antiproliferative effects reported for this formulation and its components are dose-dependent, underscoring the necessity of distinguishing therapeutic activity from potential toxicity.

*In vivo* toxicity studies are therefore an indispensable prerequisite for the responsible development and clinical translation of Vernolac. Acute oral toxicity testing provides essential information on immediate toxic effects following a single high-dose exposure, while repeated-dose (subacute/chronic) toxicity studies enable the identification of target organ toxicity, cumulative adverse effects, and safe dose limits relevant to prolonged use. Such data are particularly crucial for nutraceuticals like Vernolac, which are intended for repeated administration as supportive therapies in chronic disease settings, including cancer.

Furthermore, the availability of robust toxicological data is essential to support future *in vivo* efficacy studies, including xenograft cancer models, by enabling differentiation between adverse effects attributable to tumor burden and those arising from formulation-related toxicity [20]. In alignment with OECD guidelines for the testing of chemicals and herbal products, a structured evaluation comprising acute and repeated-dose oral toxicity studies provides a scientifically and ethically sound framework for safety assessment.

Accordingly, the present study was designed to systematically evaluate the safety of Vernolac through acute and chronic oral toxicity studies in Wistar rats. The findings of this study aim to establish critical safety parameters, including NOAEL and OELs, thereby providing foundational evidence to support the safe use of Vernolac as a nutraceutical and to inform the design of subsequent long-term and efficacy-focused investigations in cancer research.

## 2. Materials and methods

### 2.1 Poly herbal formulation

Vernolac is a commercially available polyherbal nutraceutical capsule comprising *Vernonia zeylanica* aerial parts, *Nigella sativa* seeds, *Hemidesmus indica* roots, *Smilax glabra* rhizome, and *Leucas zeylanica* aerial parts.

### 2.2 Experimental animals

Healthy albino Wistar rats of either sex, 8 - 12 weeks of age, weighing 179 ± 20 g, were purchased from Medical Research Institute, Colombo, Sri Lanka. The female rats were nulliparous and non-pregnant. The animals were maintained under standard laboratory conditions (25 °C ± 2°C, 12 h/12 h light/dark cycle, 40% - 70% relative humidity) in polypropylene cages, and had free access to standard pellet diet and water *ad libitum*. The protocol used in the study was approved by the Institute of Biology, Sri Lanka (ERC IOBSL 386/12/2024). All the animals were acclimatized for two weeks before the start of the experiment.

### 2.3 Experimental design

#### 2.3.1 Dose calculation

For the acute toxicity study, a single predefined dose of 2000 mg/kg body weight was used. Doses for the 28-day repeated dose toxicity study were selected based on the labeled human daily dose of Vernolac (1600 mg/day). Accordingly, a dose of 165 mg/kg/day was calculated as the human-equivalent dose of rats. To evaluate potential dose-dependent toxicological effects, two higher doses (330 and 660 mg/kg/day) were also selected. The individual dose administered to each rat was calculated based on its body weight.

#### 2.3.2 Acute toxicity study

The acute study was performed according to the Organisation for Economic Co-operation and Development (OECD) guideline 420 using a total of 20 healthy Wistar rats (10 male, 10 female). Animals were acclimatized for 14 days providing water and food *ad libitum.* Following acclimatizing, all the animals were fasted overnight with access to water *ad libitum*. Body weight of each animal was measured and randomly divided into two groups (n = 10 per group, five males and five females). Group 1 served as the vehicle control group and received only distilled water, while group 2 received an aqueous suspension of Vernolac at a single dose of 2000 mg/kg. Food was withheld for an additional 3 hours. The animals were then observed individually at 10 min, 30 min, 1 h, 2 h,4 h and 6 h and once daily thereafter for 14 days. During the observation, mortality, signs of toxicity, and behavioral changes (unusual vocalization, red tears in the eye, eye squinting, unusual breathing, changes in nose and cheek, ear, whisker, hunching, hypokinesia, piloerection, grooming, feces consistency, gait, nasal discharge, ocular discharge, seizure and salivation) were recorded. In addition, daily food and water consumption, and weekly body weight change were also recorded. At the end of the study period, rats were sacrificed, blood and organ samples were collected as described in 2.5.

#### 2.3.3 Repeated dose 28-day toxicity study

According to OECD Guideline 407, a repeated dose 28-day toxicity study was conducted using 40 Wistar rats. The rats were randomly divided into four groups (n = 10 per group, five males and five females). Group I served as the vehicle control group and received only distilled water, while groups II – IV received different doses of aqueous suspension of Vernolac (165, 330, and 660 mg/kg/day, respectively) for 28 consecutive days. Daily food and water consumption and weekly body weight changes were recorded throughout the study period. Further, the animals were monitored daily during the experimental period for mortality, signs of toxicity, and behavioral changes as specified in 2.3.2. On the 29^th^ day, rats were sacrificed, blood and organ samples were collected as described in 2.5.

### 2.4 Weekly body weight measurement

In both the acute and repeated-dose studies, the body weight of each rat was recorded once weekly during the 14-day acclimatization period, once prior to the initiation of dosing, weekly throughout the treatment period, and on the day of sacrifice using a calibrated sensitive balance.

### 2.5 Euthanizing, autopsy, and sample collection

Animals were sacrificed by exposure to carbon dioxide in a standard carbon dioxide euthanizing chamber. As gas levels rose to 40-50 % and animals became unconscious, the flow of gas was increased to fill the chamber more rapidly and shorten the time to death. CO_2_ flow was maintained for at least one minute after respiratory arrest. Three to five milliliters of blood samples were collected by cardiac puncture using a 16-gauge needle and a syringe from each rat under deep terminal anesthesia. The animals were then observed until all muscle activity and signs of life had been absent for at least 30 seconds. After removal from the chamber, the animals were rechecked to confirm respiratory arrest. Death was further verified by touch. If it survives, the animal was returned to the CO_2_ chamber, and death was reconfirmed.

Blood samples were collected by cardiac puncture for hematological and biochemical analysis. Approximately 1 mL of blood samples were collected into the EDTA tubes for hematological analysis. Approximately 2.5 mL of blood was collected into plain tubes for biochemical analysis and allowed to clot at room temperature, centrifuged at 2500 rpm for 10 min at room temperature to separate the serum.

Each animal underwent a detailed gross necropsy. The liver, kidney with adrenal gland, brain, heart, lung, and spleen of all animals were trimmed, and their wet weight were recorded and preserved. Moreover, relative organ weight was calculated as:

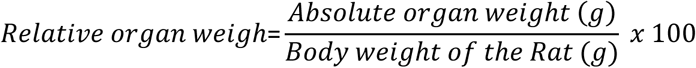

All the organs were preserved in a 4 % buffered formaldehyde solution at a 1:8 formalin-to-sample volume.

### 2.6 Hematological and biochemical analysis

White Blood Cells (WBC), neutrophils, Lymphocytes, Monocytes, Eosinophils, Basophils, Red Blood Cells (RBC), Hemoglobin, Hematocrit (HCT), Mean Corpuscular Volume (MCV), Mean Corpuscular Hemoglobin (MCH), Mean Corpuscular Hemoglobin Concentration (MCHC), Red Cell Distribution Width (RDW), and Platelets were analyzed using hematology analyzer H50, Edan Instruments Inc, China.

Biochemical analyses were performed using a calibrated automated chemistry analyzer (AS-280 E-lab biological science & technology co., ltd., China). Serum alanine aminotransferase (ALT) and aspartate aminotransferase (AST) were determined as indicators of hepatic function, while blood urea and serum creatinine were analysed to assess renal function.

### 2.7 Histopathological evaluation

Histopathological examinations were conducted on preserved organs collected from all animals in both the acute and repeated-dose toxicity studies. The organs were processed by standard paraffin embedding techniques, and tissue sections of approximately 5 μm thickness were prepared and stained with hematoxylin and eosin (H&E). Microscopic examination for treatment-related cellular alterations and morphological changes was performed using a light microscope (Olympus BX43, Japan).

### 2.8 Statistical analysis

Values from the treated groups in the acute toxicity study were compared statistically with those from the control groups using an independent-samples t-test. One-way ANOVA followed by Dunnett’s test as a post hoc analysis was used to compare treated groups with the control group in the repeated dose toxicity study. Results were expressed as mean ± SEM (n = 5); p values less than 0.05 were considered to be statistically significant. Data analysis was performed by using GraphPad Prism 8.0.1. software.

## 3 Results

### 3.1 Acute toxicity study

The acute toxicity of Vernolac was determined in accordance with OECD Guideline 420. Oral administration of Vernolac at a single dose of 2000 mg/kg was evaluated for treatment-related toxicity symptoms or mortality during the first six hours, and then daily for a period of 14 days. The results showed no change in general clinical and behavioral observations (Table 1). No significant difference (p > 0.05) was observed in body weight, food and water intake (Table 2a, 2b), relative organ weights (Table 3a, 3b), and hematological parameters (Table 4a, 4b). Only significant increases were observed in biochemical parameters, AST (p = 0.0068) and ALT (p = 0.0384), in males (Table 4b) in the treated group compared to the control group. According to the acute study findings, the LD50 for Vernolac is greater than 2000 mg/kg.

**Table 1:** General clinical and behavioral observations of the Acute Toxicity Study.

| Parameter | Control | Vernolac – 2000 mg/kg |
| --- | --- | --- |
| Body condition | Normal | Normal |
| Vocalization | Normal | Normal |
| Red tears in the eye | Not observed | Not observed |
| Eye squinting | Not observed | Not observed |
| Nose and cheek changes | Not observed | Not observed |
| Ear changes | Not observed | Not observed |
| Whisker changes | Not observed | Not observed |
| Hunching | Not observed | Not observed |
| Hypokinesia | Not observed | Not observed |
| Piloerection | Not observed | Not observed |
| Grooming | Normal | Normal |
| Breathing | Normal | Normal |
| Feces consistency | Normal | Normal |
| Gait | Normal | Normal |
| Nasal discharge | Not observed | Not observed |
| **Ocular discharge** | Not observed | Not observed |
| **Seizure** | Not observed | Not observed |
| **Salivation** | Not observed | Not observed |

**Table 2a:**
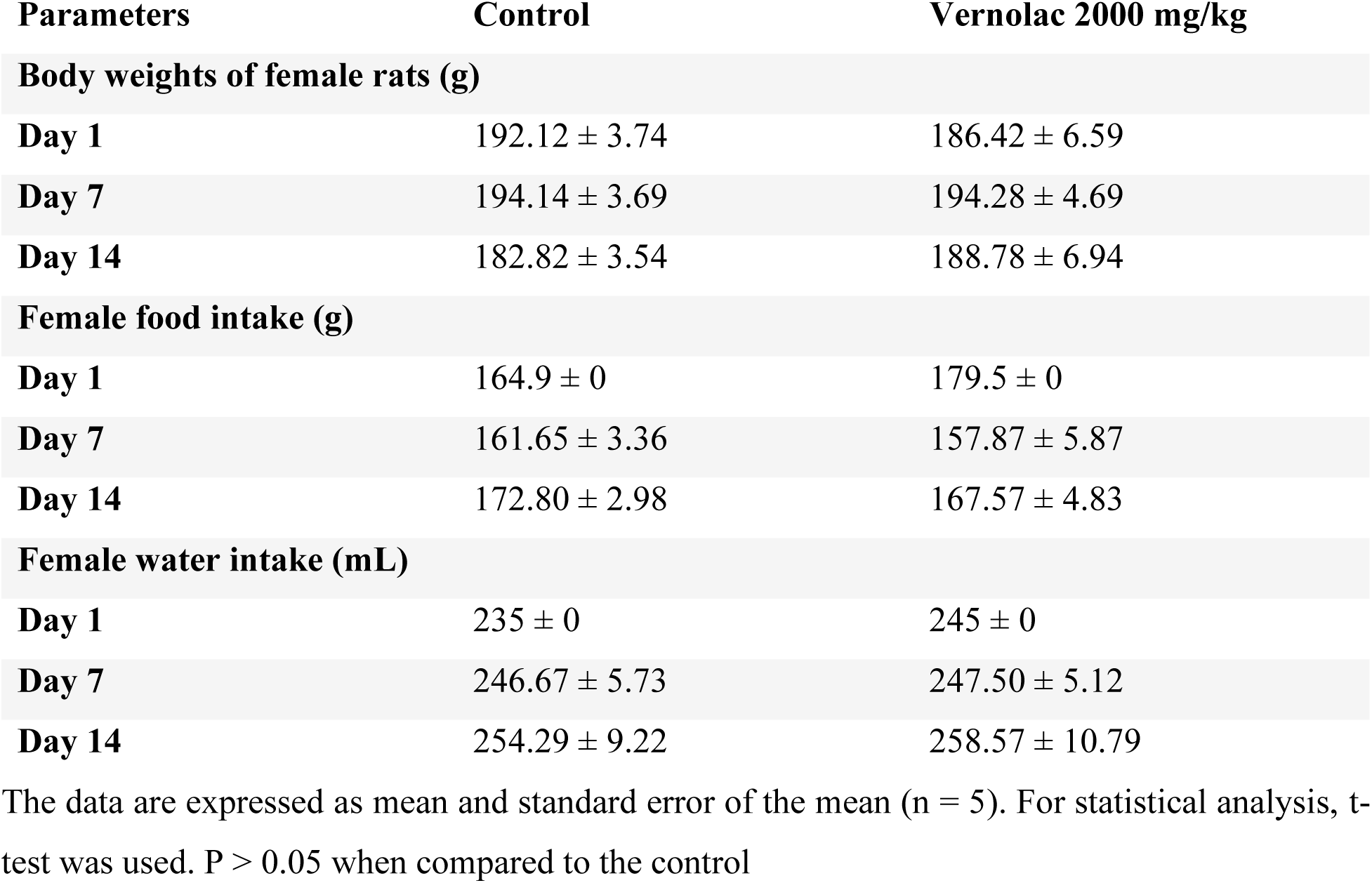
Effect of Vernolac on body weight change, food and water intake of female rats in acute toxicity study.

| **Parameters** | **Control** | **Vernolac 2000 mg/kg** |
| --- | --- | --- |
| **Body weights of female rats (g)** |  |  |
| **Day 1** | 192.12 ± 3.74 | 186.42 ± 6.59 |
| **Day 7** | 194.14 ± 3.69 | 194.28 ± 4.69 |
| **Day 14** | 182.82 ± 3.54 | 188.78 ± 6.94 |
| **Female food intake (g)** |  |  |
| **Day 1** | 164.9 ± 0 | 179.5 ± 0 |
| **Day 7** | 161.65 ± 3.36 | 157.87 ± 5.87 |
| **Day 14** | 172.80 ± 2.98 | 167.57 ± 4.83 |
| **Female water intake (mL)** |  |  |
| **Day 1** | 235 ± 0 | 245 ± 0 |
| **Day 7** | 246.67 ± 5.73 | 247.50 ± 5.12 |
| **Day 14** | 254.29 ± 9.22 | 258.57 ± 10.79 |

The data are expressed as mean and standard error of the mean (n = 5). For statistical analysis, t-test was used. P > 0.05 when compared to the control

**Table 2b:** Effect of Vernolac on body weight change, food and water intake of male rats in acute toxicity study.

| Parameters | Control | Vernolac 2000 mg/kg |
| --- | --- | --- |
| **Body weights of male rats (g)** |  |  |
| **Day 1** | 234.3 ± 6.30 | 226.06 ± 5.14 |
| **Day 7** | 248.82 ± 7.20 | 240.74 ± 4.81 |
| **Day 14** | 241.22 ± 6.31 | 233.06 ± 4.75 |
| **Male food intake (g)** |  |  |
| **Day 1** | 157.9 ± 0 | 164.2 ± 0 |
| **Day 7** | 164.60 ± 10.78 | 166.20 ± 6.86 |
| **Day 14** | 157.63 ± 3.34 | 161.71 ± 9.98 |
| **Male water intake (mL)** |  |  |
| **Day 1** | 220 ± 0 | 200.0 ± 0 |
| **Day 7** | 236.67 ± 8.43 | 250.83 ± 9.95 |
| **Day 14** | 259.29 ± 6.85 | 262.86 ± 10.17 |

The data are expressed as mean and standard error of the mean (n = 5). For statistical analysis, t-test was used. P > 0.05 when compared to the control

**Table 3a:** Effect of Vernolac on relative organ weights of female rats in acute toxicity study.

| Organ | Control | Vernolac 2000 mg/kg |
| --- | --- | --- |
| **Liver** | 3.17 ± 0.13 | 3.23 ± 0.16 |
| **Kidney + AG** | 0.33 ± 0.02 | 0.36 ± 0.03 |
| **Lung** | 0.81 ± 0.02 | 0.58 ± 0.12 |
| **Heart** | 0.35 ± 0.01 | 0.31 ± 0.03 |
| **Spleen** | 0.23 ± 0.03 | 0.25 ± 0.02 |
| **Brain** | 0.84 ± 0.05 | 0.74 ± 0.08 |

The data are expressed as mean and standard error of the mean (n = 5). For statistical analysis, t-test was used. P > 0.05 when compared to the control

**Table 3b:**
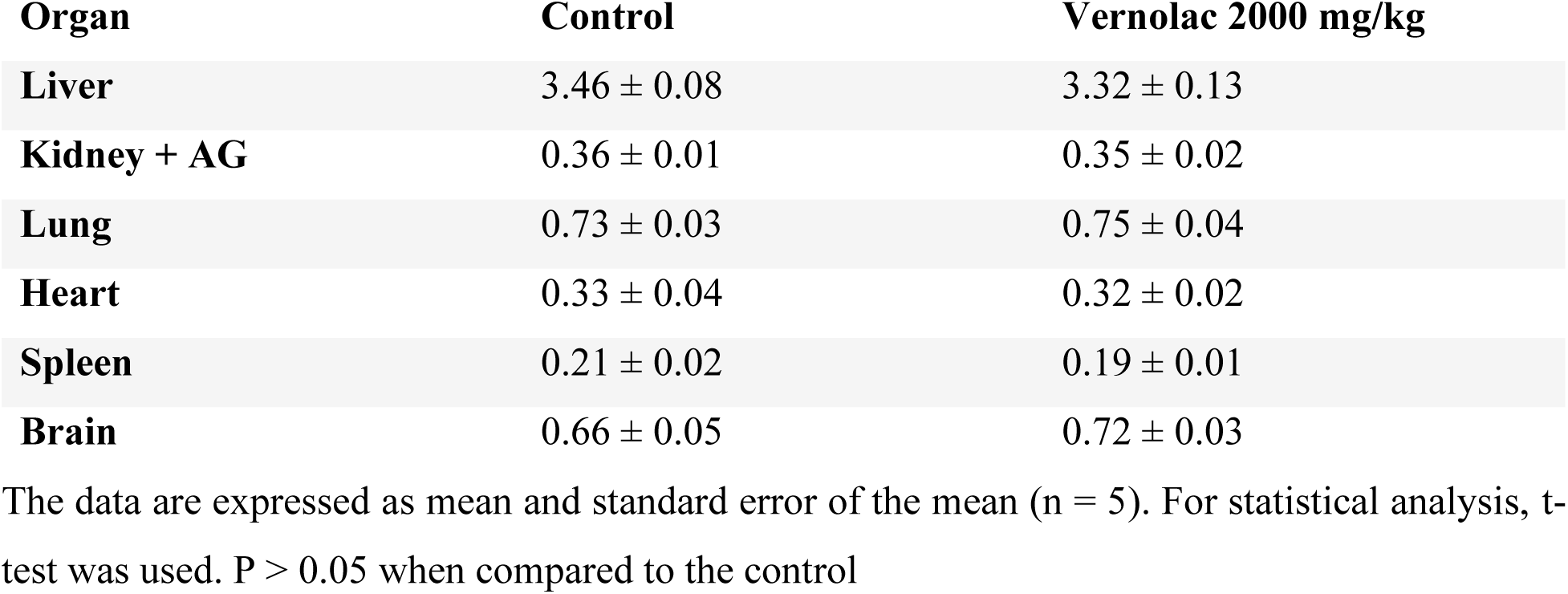
Effect of Vernolac on relative organ weights of male rats in acute toxicity study.

**Table 4a:** Effect of Vernolac on hematological and biochemical parameters of female rats in acute toxicity study.

| Parameter | Control | Vernolac 2000 mg/kg |
| --- | --- | --- |
| **Hematological parameters** |  |  |
| WBC (x10<sup>3</sup> cells/ μL) | 4.22 ± 0.61 | 6.06 ± 1.25 |
| Neutrophils (%) | 35.40 ± 1.63 | 31.60 ± 5.13 |
| Lymphocytes (%) | 59.80 ± 2.27 | 64.40 ± 5.26 |
| Monocytes (%) | 0.00 ± 0.00 | 0.60 ± 0.40 |
| Eosinophils (%) | 3.80 ± 1.11 | 3.40 ± 1.91 |
| Basophils (fL) | 0.00 ± 0.00 | 0.00 ± 0.00 |
| RBC (M/μL) | 7.13 ± 0.10 | 6.60 ± 0.43 |
| Hemoglobin (g/dl) | 13.80 ± 0.28 | 13.96 ± 0.72 |
| HCT (%) | 44.34 ± 0.35 | 42.93 ± 2.40 |
| MCV (fL) | 61.70 ± 0.49 | 67.54 ± 5.68 |
| MCH (pg) | 19.60 ± 0.18 | 21.58 ± 2.12 |
| MCHC (g/dl) | 31.62 ± 0.13 | 31.82 ± 0.40 |
| RDW (%) | 11.62 ± 0.09 | 11.83 ± 0.20 |
| **PLT (x10<sup>3</sup> cells/ μL)** | 775.6 ± 22.90 | 612.0 ± 99.32 |
| **Biochemical Parameters** |  |  |
| **AST (IU/L)** | 105.38 ± 3.87 | 206.76 ± 70.84 |
| **ALT (IU/L)** | 42.52 ± 2.10 | 65.52 ± 13.09 |
| **Blood Urea (mg/dL)** | 48.22 ± 1.72 | 42.60 ± 2.97 |
| **Serum Creatinine (mg/dL)** | 0.43 ± 0.01 | 0.45 ± 0.05 |

The data are expressed as mean and standard error of the mean (n = 5). For statistical analysis, t-test was used. P > 0.05 when compared to the control

**Table 4b:** Effect of Vernolac on hematological and biochemical parameters of male rats in acute toxicity study.

| **Parameter** | **Control** | **Vernolac 2000 mg/kg** |
| --- | --- | --- |
| **Hematological parameters** |  |  |
| **WBC (x10<sup>3</sup> cells/ μL)** | 6.94 ± 0.79 | 7.48 ± 0.38 |
| **Neutrophils (%)** | 29 ± 2.47 | 30.4 ± 2.42 |
| **Lymphocytes (%)** | 70.4 ± 2.36 | 67.8 ± 2.42 |
| **Monocytes (%)** | 0.2 ± 0.20 | 0.8 ± 0.20 |
| **Eosinophils (%)** | 0.4 ± 0.24 | 1 ± 0.32 |
| **Basophils (fL)** | 0 ± 0 | 0 ± 0 |
| **RBC (M/μL)** | 7.858 ± 0.14 | 8.12 ± 0.10 |
| **Hemoglobin (g/dl)** | 15.2 ± 0.37 | 15.26 ± 0.17 |
| **HCT (%)** | 48.02 ± 1.16 | 49.02 ± 0.78 |
| **MCV (fL)** | 61.08 ± 0.45 | 60.36 ± 0.48 |
| **MCH (pg)** | 19.34 ± 0.20 | 18.8 ± 0.11 |
| **MCHC (g/dl)** | 31.66 ± 0.16 | 31.14 ± 0.22 |
| **RDW (%)** | 13.1 ± 0.14 | 13.1 ± 0.26 |
| **PLT (x10<sup>3</sup> cells/ μL)** | 814.8 ± 9.12 | 761.0 ± 21.93 |
| **Biochemical Parameters** |  |  |
| **AST (IU/L)** | 109.9 ± 5.60 | 148.82 ± 8.54** |
| **ALT (IU/L)** | 50.78 ± 2.48 | 58.16 ± 1.29* |
| **Blood Urea (mg/dL)** | 46.58 ± 1.55 | 49.64 ± 2.08 |
| **Serum Creatinine (mg/dL)** | 0.394 ± 0.01 | 0.384 ± 0.02 |

The data are expressed as mean and standard error of the mean (n = 5). For statistical analysis, t-test was used. \* Represents significance of significance of p<0.05, \*\* represents significance of significance of p<0.01

### 3.2 Repeated dose 28-day oral toxicity study

The daily oral administration of Vernolac at doses of 165, 330, and 660 mg/kg/day for 28 consecutive days did not result in any adverse symptoms, mortality, or abnormal clinical signs (Table 5).

**Table 5:**
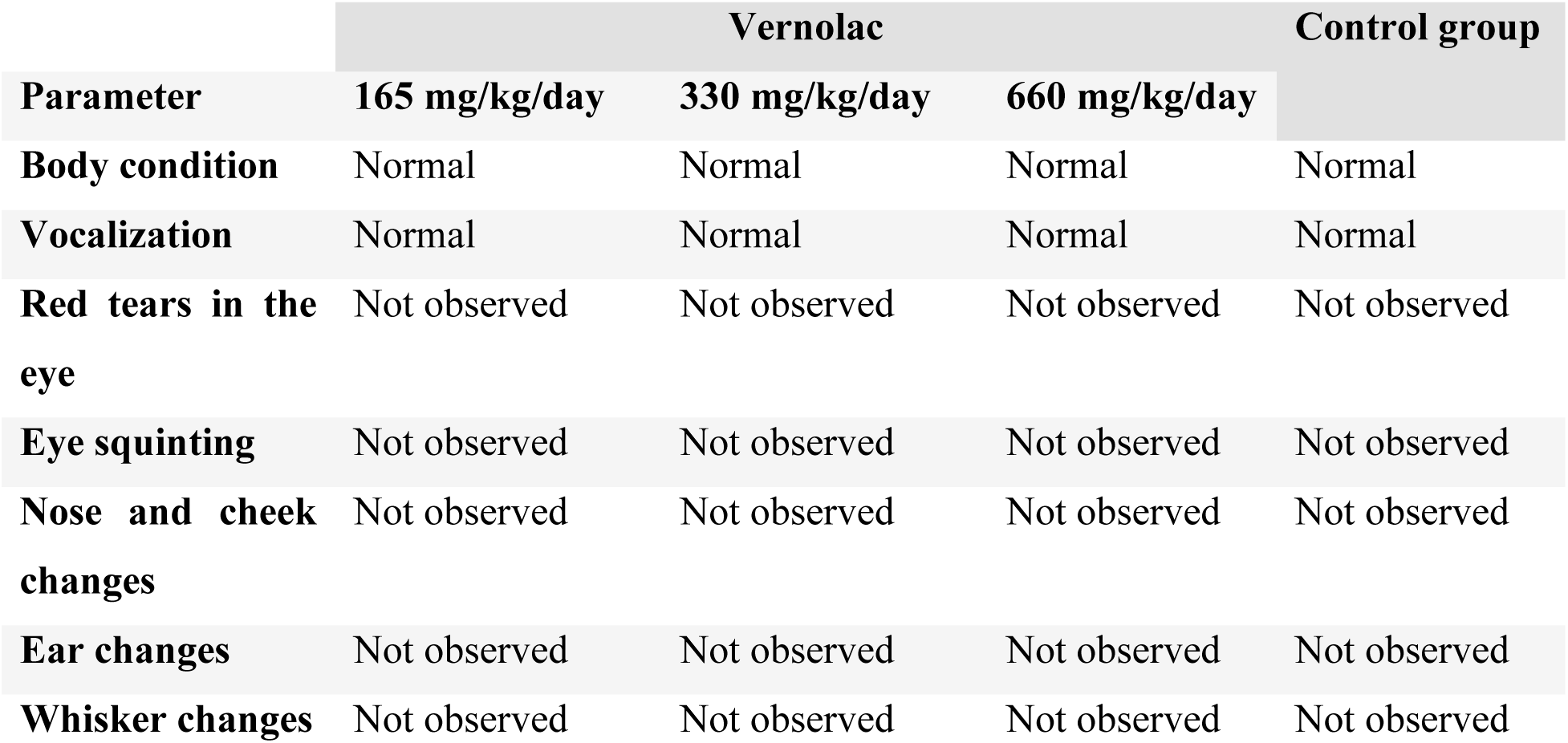
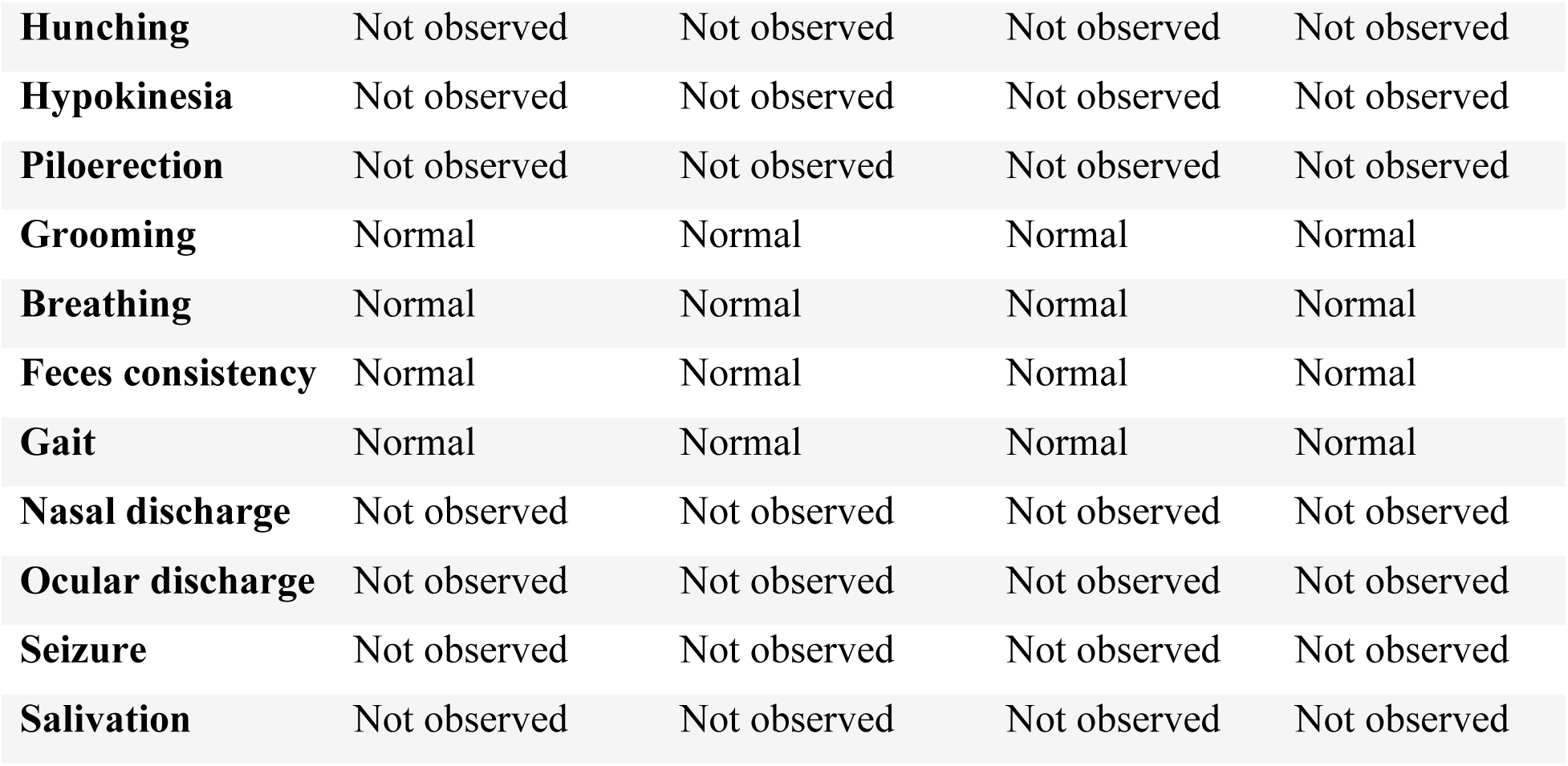
General clinical and behavioral observations of the 28-day repeated dose toxicity study for the control and Vernolac-treated groups.

| **Parameter** | **Vernolac** |  |  | **Control group** |
| --- | --- | --- | --- | --- |
|  | **165 mg/kg/day** | **330 mg/kg/day** | **660 mg/kg/day** |  |
| **Body condition** | Normal | Normal | Normal | Normal |
| **Vocalization** | Normal | Normal | Normal | Normal |
| **Red tears in the eye** | Not observed | Not observed | Not observed | Not observed |
| **Eye squinting** | Not observed | Not observed | Not observed | Not observed |
| **Nose and cheek changes** | Not observed | Not observed | Not observed | Not observed |
| **Ear changes** | Not observed | Not observed | Not observed | Not observed |
| **Whisker changes** | Not observed | Not observed | Not observed | Not observed |
| **Hunching** | Not observed | Not observed | Not observed | Not observed |
| **Hypokinesia** | Not observed | Not observed | Not observed | Not observed |
| **Piloerection** | Not observed | Not observed | Not observed | Not observed |
| **Grooming** | Normal | Normal | Normal | Normal |
| **Breathing** | Normal | Normal | Normal | Normal |
| **Feces consistency** | Normal | Normal | Normal | Normal |
| **Gait** | Normal | Normal | Normal | Normal |
| **Nasal discharge** | Not observed | Not observed | Not observed | Not observed |
| **Ocular discharge** | Not observed | Not observed | Not observed | Not observed |
| **Seizure** | Not observed | Not observed | Not observed | Not observed |
| **Salivation** | Not observed | Not observed | Not observed | Not observed |

#### 3.2.1 Effect on body weight

Body weights of male and female rats in the control and Vernolac-treated groups (165, 330, and 660 mg/kg/day) were monitored throughout the treatment period. Minor fluctuations in body weight were observed in all groups; however, no statistically significant differences (p > 0.05) were detected between control and treated females at any dose level (Table 6a). In male rats (Table 6b), a statistically significant increase in final body weight was observed only in the 165 mg/kg/day group compared with the control (p < 0.01). No significant differences in body weight were observed at the higher dose levels (330 and 660 mg/kg/day).

**Table 6a:**
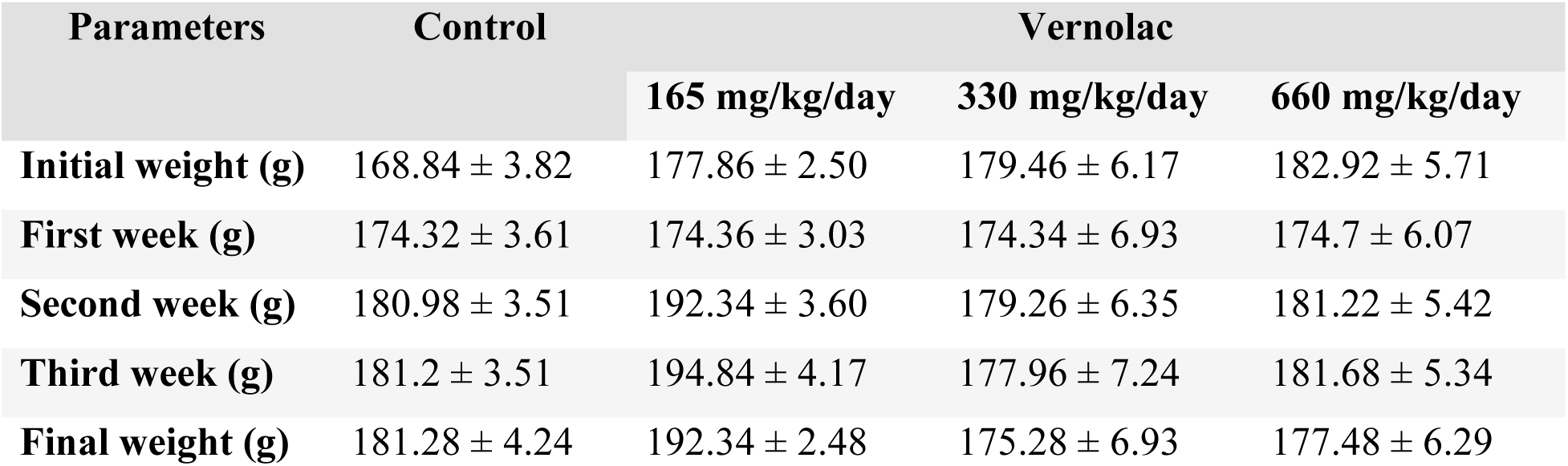
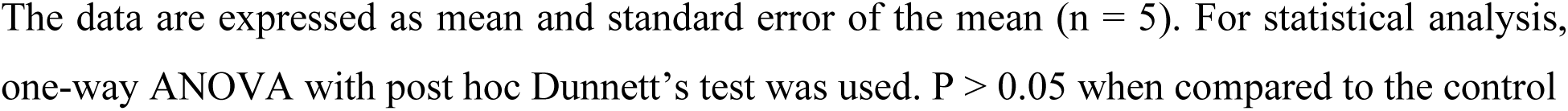
Body weights of female Wistar rats treated with Vernolac and the control group in repeated dose 28-day oral toxicity study.

**Table 6b:** Body weights of male Wistar rats treated with Vernolac and the control group in repeated dose 28-day oral toxicity study.

| Parameters | Control | Vernolac |  |  |
| --- | --- | --- | --- | --- |
|  |  | 165 mg/kg/day | 330 mg/kg/day | 660 mg/kg/day |
| **Initial weight (g)** | 191.94 ± 4.93 | 205.2 ± 3.49 | 173.58 ± 3.34 | 193.52 ± 5.43 |
| **First week (g)** | 193.5 ± 4.90 | 205.22 ± 3.82 | 186.08 ± 2.43 | 194.86 ± 5.81 |
| **Second week (g)** | 199 ± 4.64 | 216.24 ± 3.99 | 198.46 ± 2.70 | 193.68 ± 7.99 |
| **Third week (g)** | 200.98 ± 3.71 | 211.96 ± 5.28 | 195.46 ± 2.46 | 197.85 ± 10.30 |
| **Final weight (g)** | 193.48 ± 4.27 | 206.14 ± 5.01** | 190.4 ± 1.61 | 201.85 ± 10.58 |

The data are expressed as mean and standard error of the mean (n = 5). For statistical analysis, one-way ANOVA with post hoc Dunnett's test was used. \*\* represents significance of p<0.01

#### 3.2.2 Effect on food and water consumption

Daily food and water intake in female and male rats treated with Vernolac at doses of 165, 330, and 660 mg/kg/day for 28 days did not differ significantly (p > 0.05) from those of the respective control groups throughout the treatment period (Table 7a and 7b).

**Table 7a:**
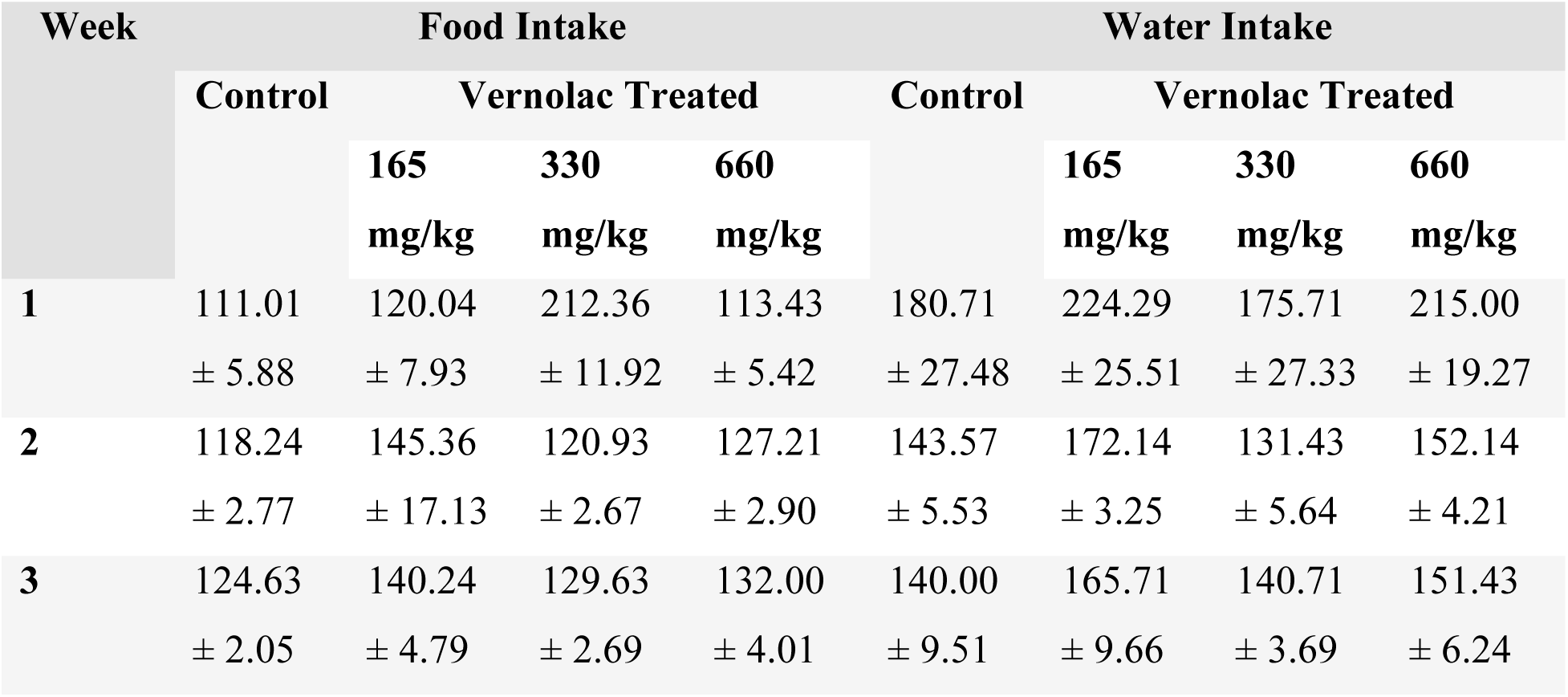
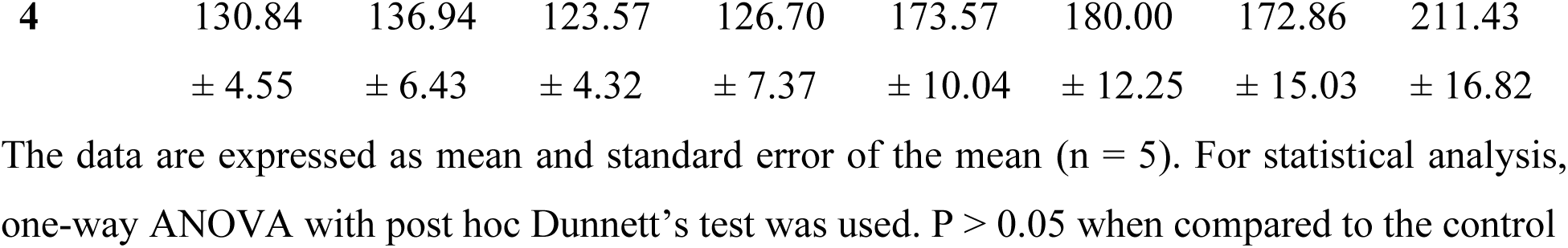
Food and water intake of female Wistar rats in 28-day repeated dose toxicity study.

**Table 7b:**
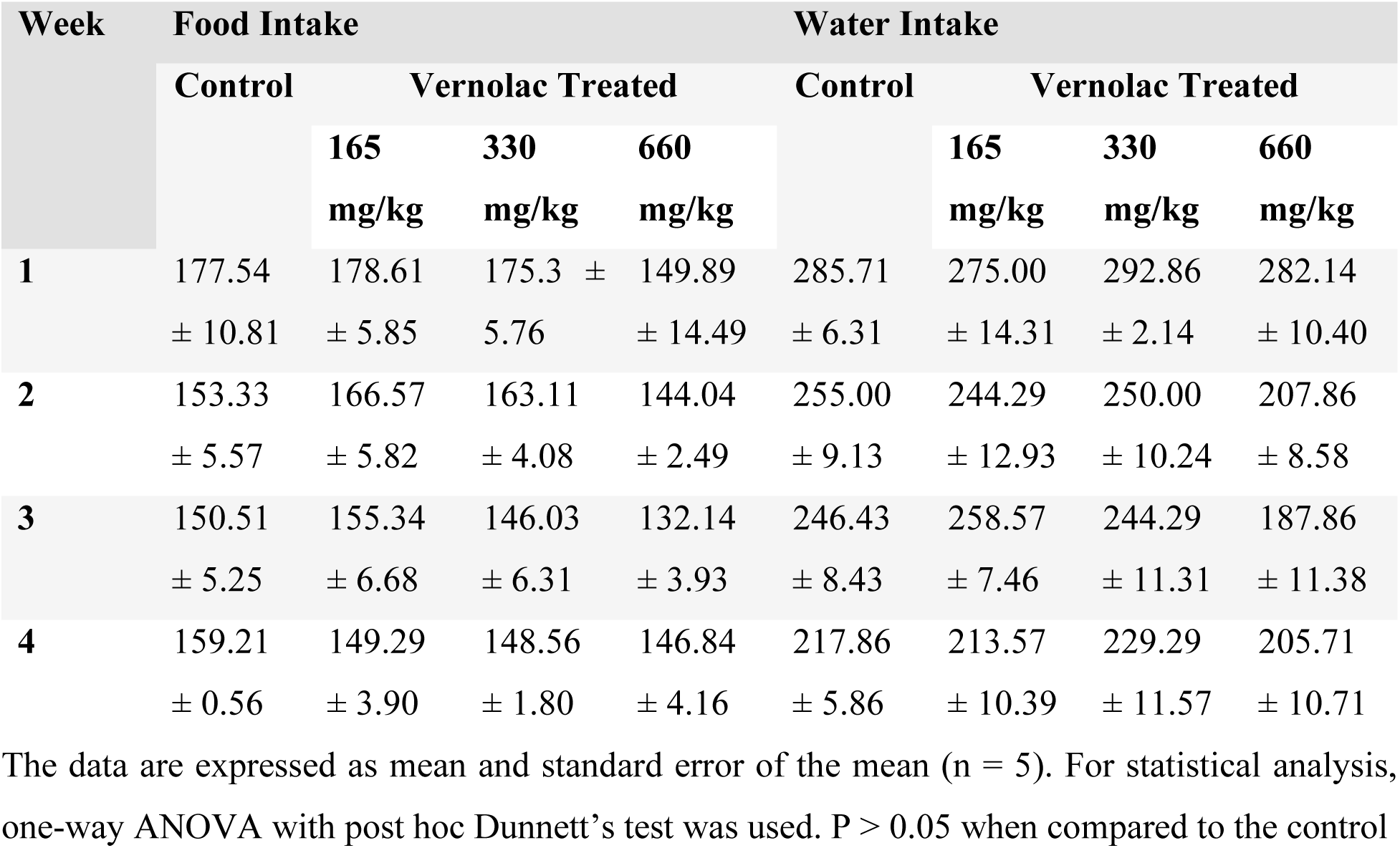
Food and water intake of male Wistar rats in 28-day repeated dose toxicity study.

#### 3.2.3 Effect on relative organ weight

No statistically significant differences (p > 0.05) were observed in the relative organ weights of female and male rats in the repeated-dose toxicity study when compared with the respective control groups (Tables 8a and 8b).

**Table 8a:** Relative organ weights of female Wistar rats in 28-day repeated dose toxicity study.

| Organ | Control | Vernolac |  |  |
| --- | --- | --- | --- | --- |
|  |  | 165 mg/kg | 330 mg/kg | 660 mg/kg |
| **Liver** | 3.22 ± 0.09 | 3.13 ± 0.12 | 3.28 ± 0.12 | 3.05 ± 0.12 |
| **Kidney + AG** | 0.27 ± 0.02 | 0.30 ± 0.01 | 0.30 ± 0.01 | 0.30 ± 0.01 |
| **Lung** | 0.79 ± 0.02 | 0.65 ± 0.04 | 0.80 ± 0.02 | 0.74 ± 0.03 |
| **Heart** | 0.39 ± 0.01 | 0.43 ± 0.02 | 0.39 ± 0.02 | 0.33 ± 0.03 |
| **Spleen** | 0.21 ± 0.01 | 0.20 ± 0.02 | 0.17 ± 0.01 | 0.23 ± 0.01 |
| **Brain** | 0.88 ± 0.02 | 0.82 ± 0.03 | 0.91 ± 0.04 | 0.91 ± 0.03 |

The data are expressed as mean and standard error of the mean (n = 5). For statistical analysis, one-way ANOVA with post hoc Dunnett's test was used. P > 0.05 when compared to the control

**Table 8b:** Relative organ weights of male Wistar rats in 28-day repeated dose toxicity study.

| Organ | Control | Vernolac |  |  |
| --- | --- | --- | --- | --- |
|  |  | 165 mg/kg | 330 mg/kg | 660 mg/kg |
| **Liver** | 3.28 ± 0.07 | 3.21 ± 0.16 | 3.51 ± 0.12 | 3.19 ± 0.13 |
| **Kidney + AG** | 0.33 ± 0.01 | 0.34 ± 0.01 | 0.36 ± 0.02 | 0.33 ± 0.02 |
| **Lung** | 0.52 ± 0.14 | 0.71 ± 0.04 | 0.80 ± 0.01 | 0.75 ± 0.06 |
| **Heart** | 0.31 ± 0.03 | 0.36 ± 0.01 | 0.31 ± 0.03 | 0.36 ± 0.01 |
| **Spleen** | 0.18 ± 0.01 | 0.17 ± 0.02 | 0.17 ± 0.01 | 0.17 ± 0.01 |
| **Brain** | 0.82 ± 0.03 | 0.77 ± 0.02 | 0.75 ± 0.03 | 0.79 ± 0.05 |

The data are expressed as mean and standard error of the mean (n = 5). For statistical analysis, one-way ANOVA with post hoc Dunnett's test was used. P > 0.05 when compared to the control

#### 3.2.4 Effect on hematological and biochemical blood parameters

In female rats (Table 9a), no statistically significant differences were observed in hematological and biochemical parameters compared with the control group, except for MCH and ALT, which showed statistically significant changes only at the highest dose level. In male rats (Table 9b), no significant differences were observed in WBC, neutrophils, lymphocytes, monocytes, basophils, RBC, or MCHC when compared with the control group. A marked and statistically significant reduction in eosinophil counts was observed in all Vernolac-treated male groups, with complete depletion noted at the highest dose level. Variations in hemoglobin, HCT, MCV, MCH, RDW, and platelet counts were observed; however, all values remained within established physiological reference ranges.

**Table 9a:**
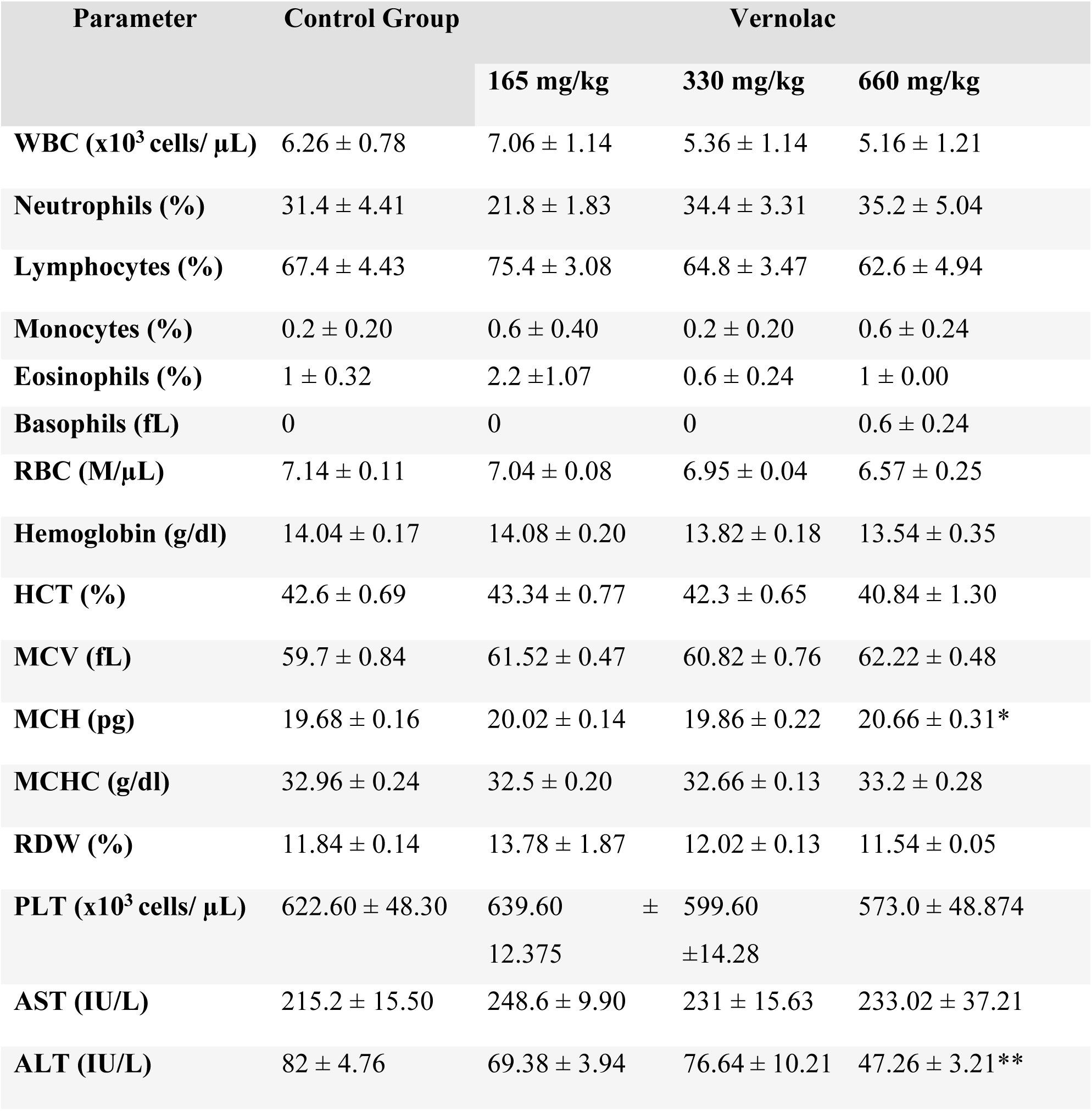
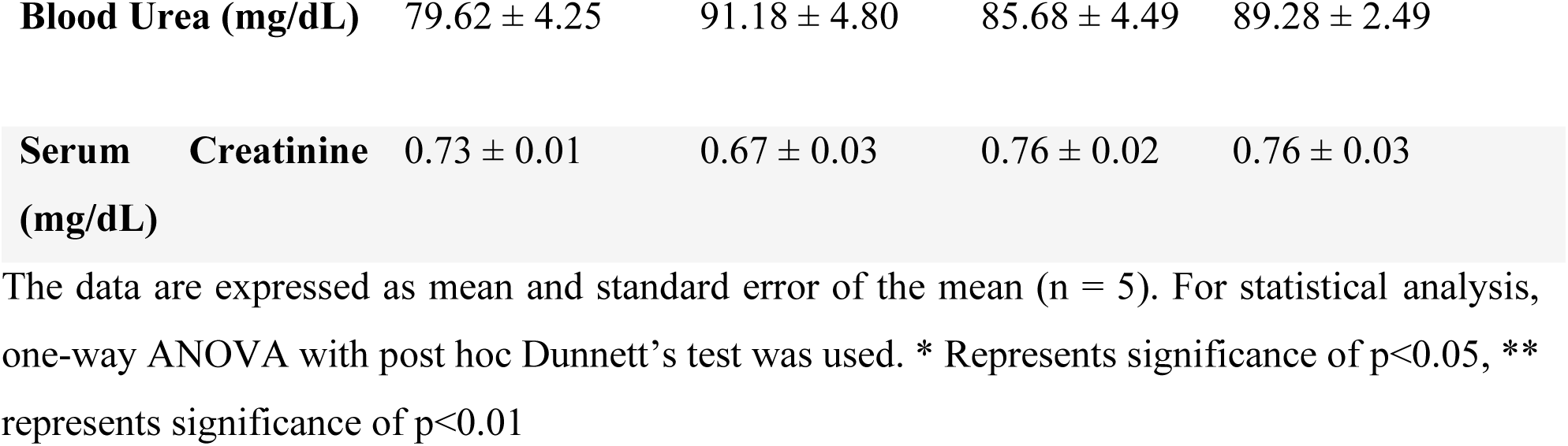
Effect of Vernolac on hematological and biochemical parameters on female Wistar rats in 28-day repeated dose toxicity study.

**Table 9b:**
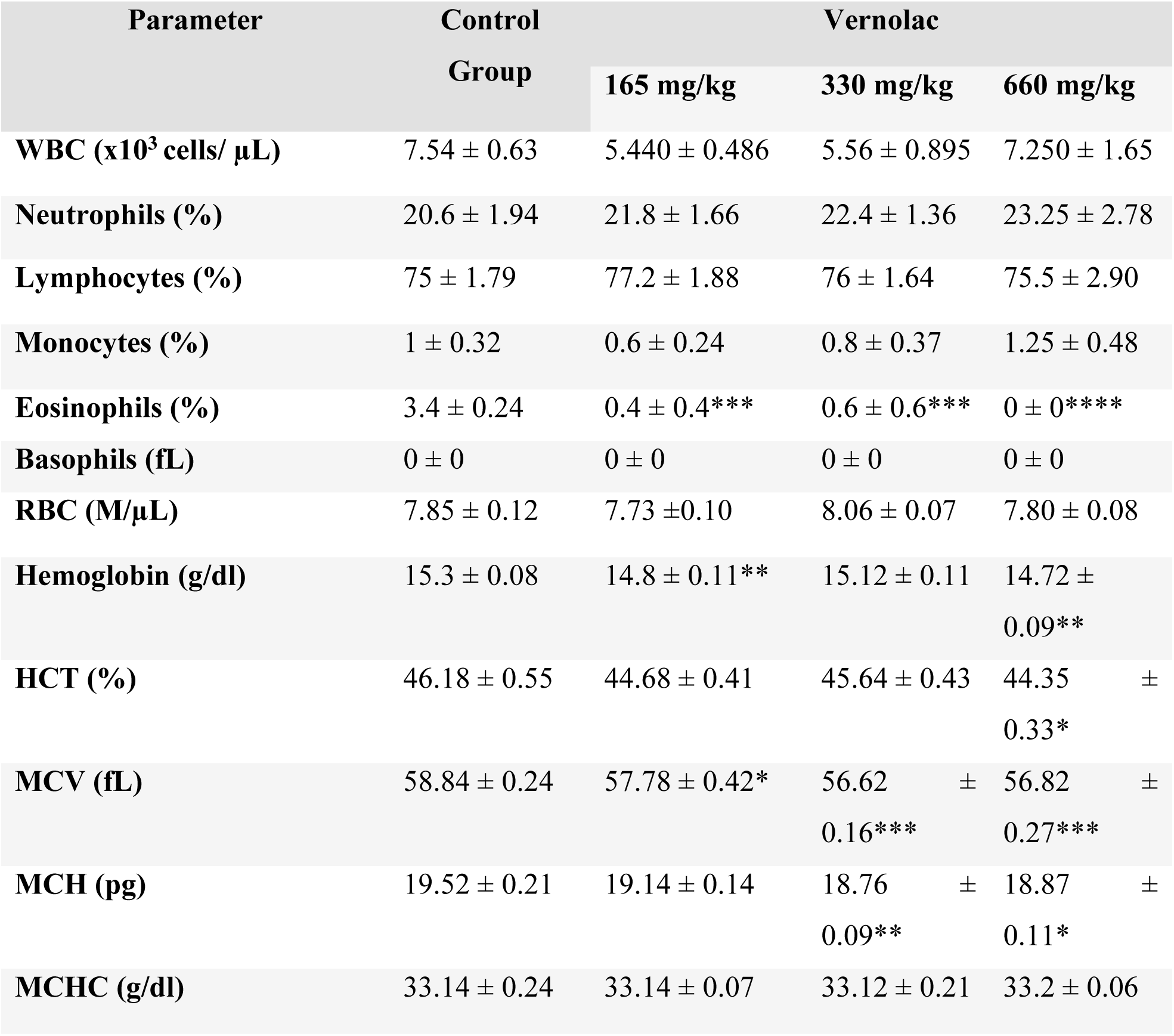
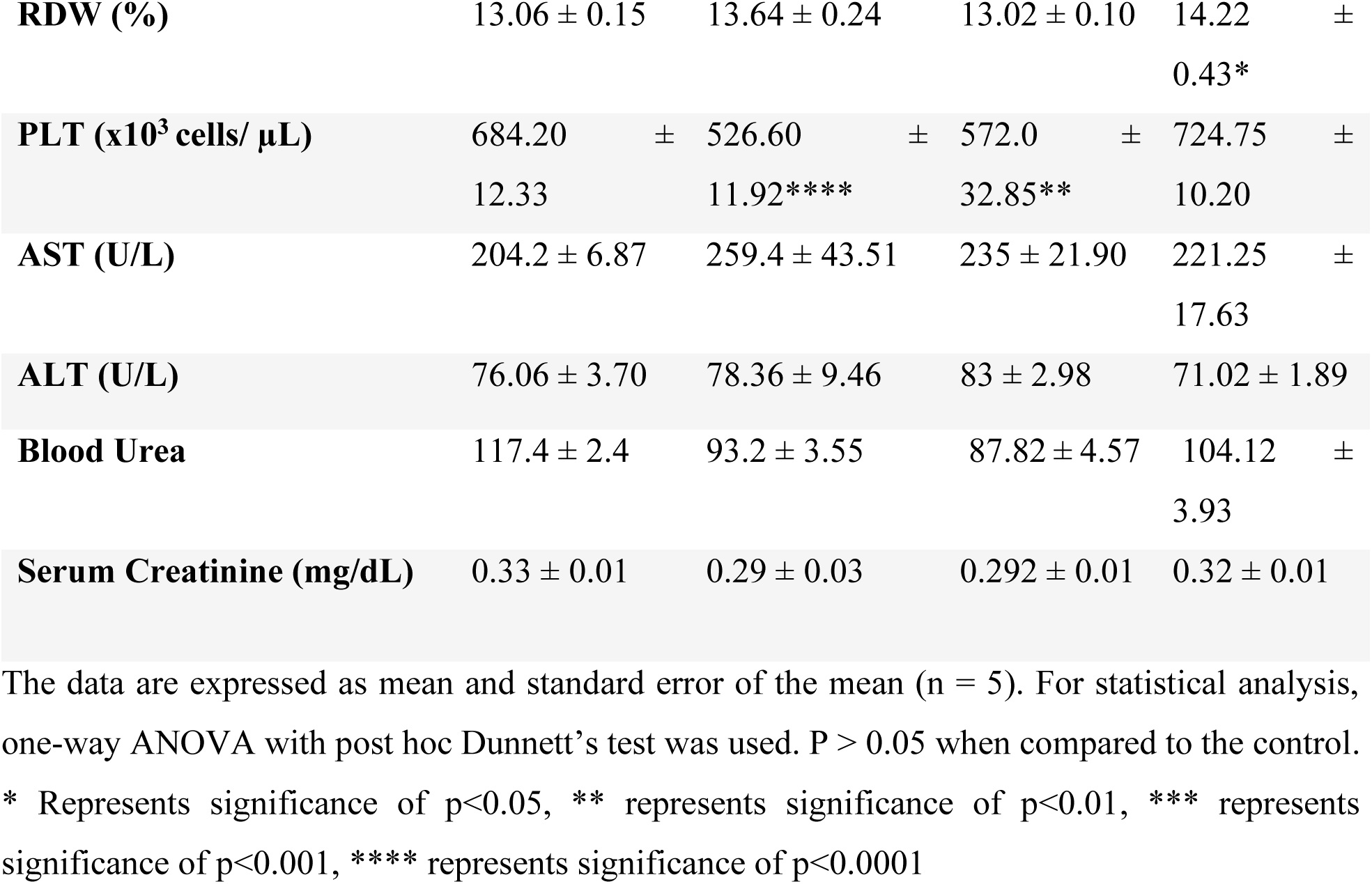
Effect of Vernolac on hematological and biochemical parameters on male Wistar rats in 28-days repeated dose toxicity study.

| Parameter | Control Group | Vernolac |  |  |
| --- | --- | --- | --- | --- |
|  |  | 165 mg/kg | 330 mg/kg | 660 mg/kg |
| **WBC (x10<sup>3</sup> cells/ µL)** | 7.54 ± 0.63 | 5.440 ± 0.486 | 5.56 ± 0.895 | 7.250 ± 1.65 |
| **Neutrophils (%)** | 20.6 ± 1.94 | 21.8 ± 1.66 | 22.4 ± 1.36 | 23.25 ± 2.78 |
| **Lymphocytes (%)** | 75 ± 1.79 | 77.2 ± 1.88 | 76 ± 1.64 | 75.5 ± 2.90 |
| **Monocytes (%)** | 1 ± 0.32 | 0.6 ± 0.24 | 0.8 ± 0.37 | 1.25 ± 0.48 |
| **Eosinophils (%)** | 3.4 ± 0.24 | 0.4 ± 0.4*** | 0.6 ± 0.6*** | 0 ± 0**** |
| **Basophils (fL)** | 0 ± 0 | 0 ± 0 | 0 ± 0 | 0 ± 0 |
| **RBC (M/µL)** | 7.85 ± 0.12 | 7.73 ± 0.10 | 8.06 ± 0.07 | 7.80 ± 0.08 |
| **Hemoglobin (g/dl)** | 15.3 ± 0.08 | 14.8 ± 0.11** | 15.12 ± 0.11 | 14.72 ± 0.09** |
| **HCT (%)** | 46.18 ± 0.55 | 44.68 ± 0.41 | 45.64 ± 0.43 | 44.35 ± 0.33* |
| **MCV (fL)** | 58.84 ± 0.24 | 57.78 ± 0.42* | 56.62 ± 0.16*** | 56.82 ± 0.27*** |
| **MCH (pg)** | 19.52 ± 0.21 | 19.14 ± 0.14 | 18.76 ± 0.09** | 18.87 ± 0.11* |
| **MCHC (g/dl)** | 33.14 ± 0.24 | 33.14 ± 0.07 | 33.12 ± 0.21 | 33.2 ± 0.06 |
| **RDW (%)** | 13.06 ± 0.15 | 13.64 ± 0.24 | 13.02 ± 0.10 | 14.22 ± 0.43* |
| **PLT (x10<sup>3</sup> cells/ µL)** | 684.20 ± 12.33 | 526.60 ± 11.92**** | 572.0 ± 32.85** | 724.75 ± 10.20 |
| **AST (U/L)** | 204.2 ± 6.87 | 259.4 ± 43.51 | 235 ± 21.90 | 221.25 ± 17.63 |
| **ALT (U/L)** | 76.06 ± 3.70 | 78.36 ± 9.46 | 83 ± 2.98 | 71.02 ± 1.89 |
| **Blood Urea** | 117.4 ± 2.4 | 93.2 ± 3.55 | 87.82 ± 4.57 | 104.12 ± 3.93 |
| **Serum Creatinine (mg/dL)** | 0.33 ± 0.01 | 0.29 ± 0.03 | 0.292 ± 0.01 | 0.32 ± 0.01 |

The data are expressed as mean and standard error of the mean (n = 5). For statistical analysis, one-way ANOVA with post hoc Dunnett's test was used. P > 0.05 when compared to the control. \* Represents significance of p<0.05, \*\* represents significance of p<0.01, \*\*\* represents significance of p<0.001, \*\*\*\* represents significance of p<0.0001

### 3.3 Histological examinations

Sections of liver, spleen, kidney, lung, heart, and brain were used for histopathological analysis in both the acute and repeated dose toxicity studies.

#### 3.3.1 Liver

In the acute (Fig. 1) and 28-days repeated dose studies (Fig. 2A, and B), the histopathological examination of liver sections of all male and female rats showed preserved hepatic architecture (Fig. 1).

**Fig. 1.**
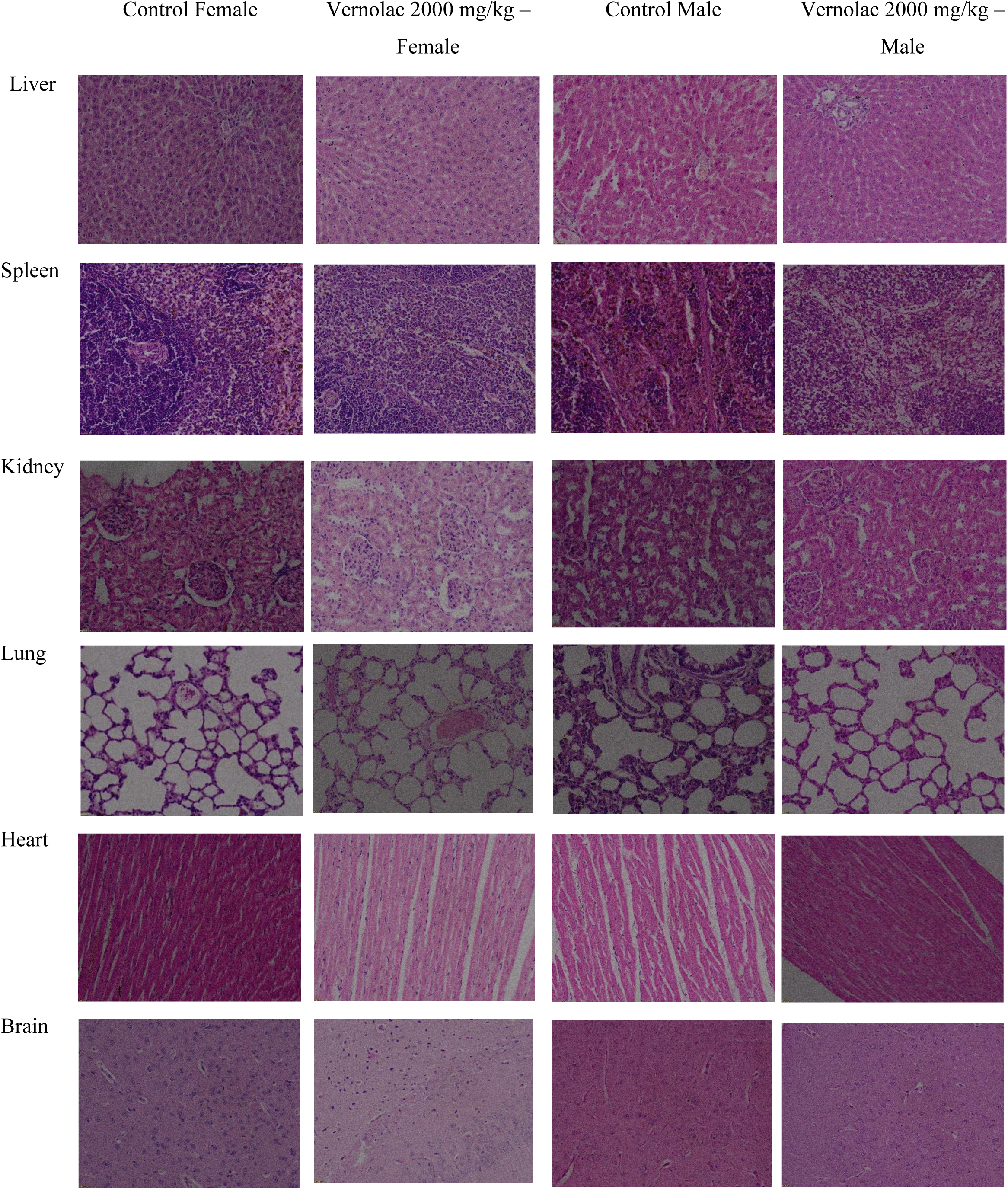
Histopathological images of organs (liver, spleen, kidney, lung, heart, and brain) in the acute toxicity study at 400x. Liver with preserved hepatic architecture. Spleen with standard architecture, including red and white pulp. Kidney with preserved renal architecture. Lung, heart, and brain with their preserved architecture.

**Fig. 2.**
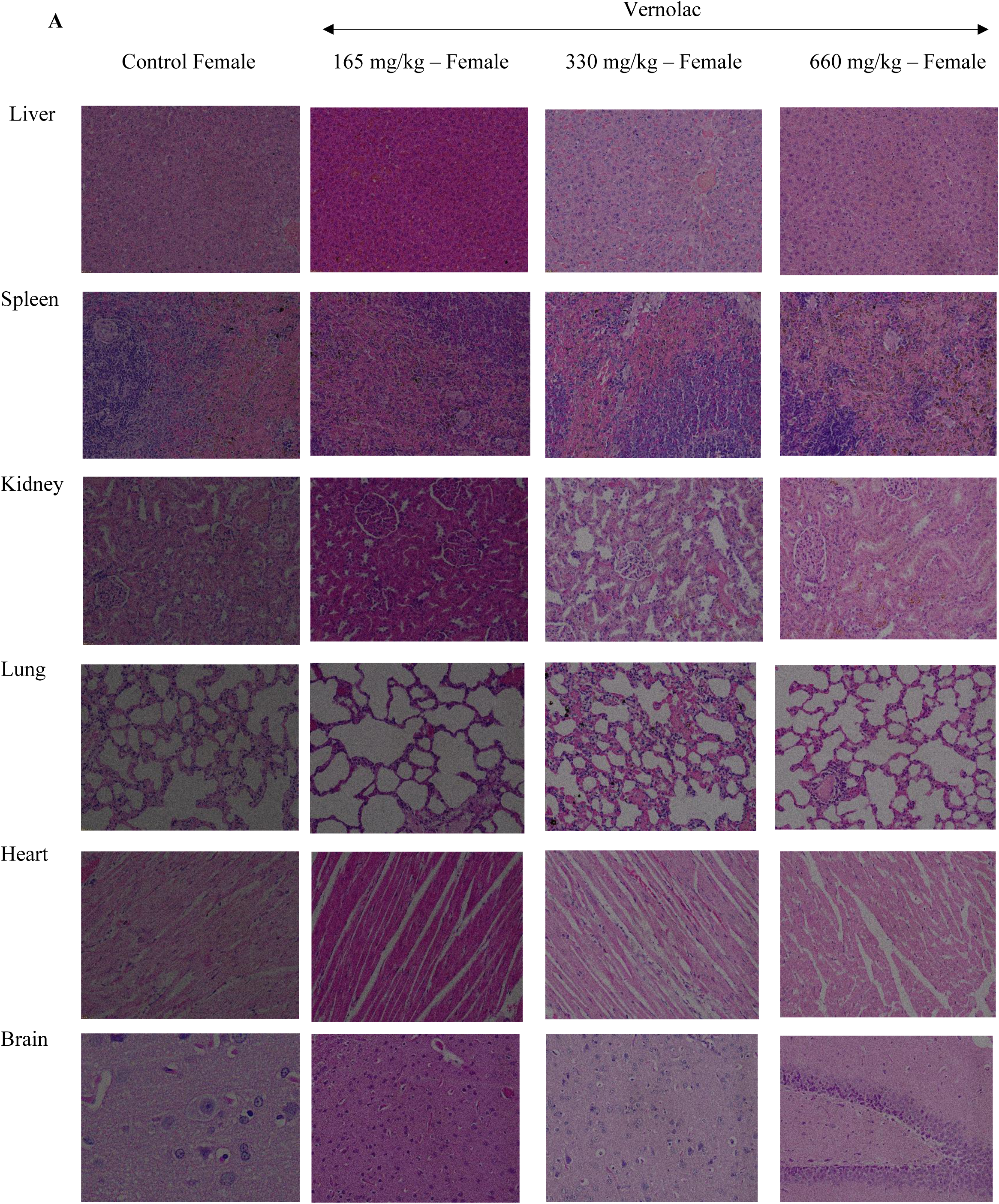
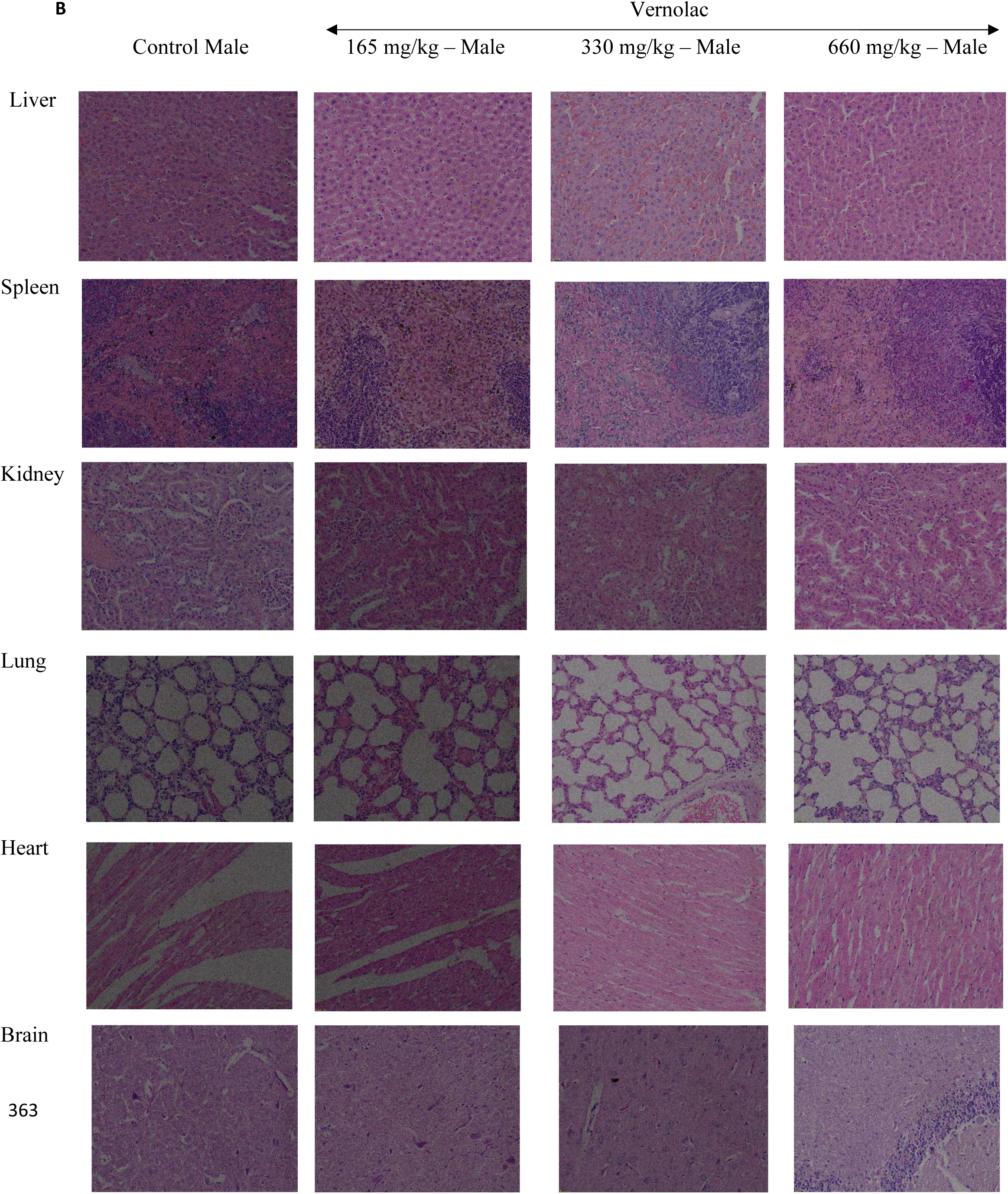
**A.** Histopathological images of organs (liver, spleen, kidney, lung, heart, and brain) of female rats in 28 days repeated dose toxicity study treated with Vernolac at doses 165 mg/kg, 330 mg/kg, 660 mg/kg/day and the control group at 400x. **B** Histopathological images of organs (liver, spleen, kidney, lung, heart, and brain) of male rats in 28 days repeated dose toxicity study treated with Vernolac at doses 165 mg/kg, 330 mg/kg, 660 mg/kg/day, and the control group at 400x. In both A and B, hepatic architecture was preserved in liver sections. Spleen with standard architecture, including red and white pulp. Kidney with preserved renal architecture. Lung, heart, and brain with their standard preserved architecture.

#### 3.3.2 Spleen

In both the acute (Fig. 1) and 28-days repeated dose studies (Fig. 2A, and B), all rats in the control and treated groups demonstrated preserved spleen architecture, with intact red pulp and white pulp compartments.

#### 3.3.3 Kidney

In both the acute (Fig. 1) and 28-days repeated dose studies (Fig. 2A, and B), histopathological examination of kidney tissues demonstrated preserved renal architecture. Glomeruli, renal tubules, and renal blood vessels appeared viable and normal in all rats in the treated and control groups. The interstitium was normal with no inflammation or fibrosis observed across all rats.

#### 3.3.4 Lung

In both the acute (Fig. 1) and 28-days repeated dose studies (Fig. 2A, and B), microscopic examination of lung sections revealed no significant differences between control and Vernolac-treated rats.

#### 3.3.5 Heart

In both the acute (Fig. 1) and 28-days repeated dose studies (Fig. 2A, and B), microscopic examination of heart sections revealed no significant differences between control and Vernolac-treated rats.

#### 3.3.6 Brain

In both the (Fig. 1) and 28-days repeated dose studies (Fig. 2A, and B), microscopic examinations of brain sections revealed no significant differences between control and Vernolac-treated rats.

## 4 Discussion

Vernolac is a commercially available polyherbal formulation used in cancer therapy, comprising the aerial parts of *Vernonia zeylanica*, seeds of *Nigella sativa*, roots of *Hemidesmus indicus*, rhizomes of *Smilax glabra*, and the aerial parts of *Leucas zeylanica*. Several studies have evaluated the anticancer potential of Vernolac as a formulation, as well as that of its individual plant components and selected isolated phytochemicals. A recently published study reports the promising therapeutic potential of Vernolac targeting cancer stem cells [12]. Vernolac supercritical CO_2_ extract showed potent anticancer activity against cancer stem-like (CSCL) NTERA-2 cl. D1 Cells, with an IC_50_ of 41.12 µg/mL at 48 h. Fluorescence microscopy and caspase 3/7 assay confirmed apoptosis induction, along with increased ROS levels, inhibited cell migration, and regulation of key genes (↑TP53, ↓Survivin, ↓mTOR) [12]. A network pharmacology-based study indicated that Vernolac may exert anticancer effects through mechanisms such as induction of apoptosis, immune modulation, anti-inflammatory and anti-proliferative activities, as well as chemoradiosensitizing properties. It was also reported that Vernolac may exhibit chemoradioprotective potential by alleviating therapy-induced toxicity, supporting its promise as an adjunct to conventional cancer treatments [19]. In a separate in vitro study, the cytotoxic effects of a decoction composed of *Nigella sativa* seeds, *Hemidesmus indicus* roots, and *Smilax glabra* rhizomes, as well as the corresponding individual plant extracts, were evaluated using the human hepatoma HepG2 cell line. The combined decoction demonstrated a pronounced, dose-dependent cytotoxic effect on HepG2 cells. Among the individual extracts, cytotoxic potency was highest for *N. sativa*, followed by *H. indicus* and *S. glabra* [13]. Another study confirmed that this standardized polyherbal decoction induces apoptosis in HepG2 cells through multiple mechanisms, including DNA fragmentation and characteristic apoptotic morphological changes.

The treatment was shown to up-regulate the expression of the pro-apoptotic gene *Bax* while down-regulating the anti-apoptotic gene *Bcl-2*, along with time- and dose-dependent activation of caspase-3 and caspase-9 [14]. The anti-inflammatory properties of this polyherbal decoction have been proposed to contribute to its reported anti-hepatocarcinogenic activity. In vivo studies demonstrated that the decoction significantly reduced carrageenan-induced paw edema and suppressed hepatic NF-κB activation through inhibition of IKKα/β in rats. In addition, the decoction was shown to inhibit nitric oxide production, reduce leukocyte migration, and stabilize cell membranes, further supporting its anti-inflammatory potential [15]. Furthermore, several studies have proven the anticancer activity of the active compounds isolated from the above plants [16, 17, 18].

The present study evaluated the *in vivo* safety and toxicity of Vernolac. Toxicological studies using animal models are essential for predicting the potential effects of new substances on human health, as they help identify organ-specific effects and establish safe exposure levels [21]. Acute and 28-day repeated dose oral toxicity studies in Wistar rats were conducted to assess the safety profile of Vernolac.

The oral acute toxicity of Vernolac was assessed following a single administration at a dose of 2000 mg/kg. Animals were observed for a 14-day period, during which no mortality, treatment-related behavioral changes, or visible signs of toxicity were detected. No statistically significant differences were observed in body weight gain, food and water consumption, or relative organ weights in either male or female rats when compared with the respective control groups. In female Wistar rats, no significant alterations were observed in hematological or biochemical parameters. Similarly, hematological parameters in male rats did not differ significantly from those of the control group. However, serum AST and alanine ALT activities were significantly elevated in male rats compared with controls. No association was identified between the altered biochemical parameters and the histopathological findings in liver sections. Similarly, a long-term study of 16 months conducted by Iddamaldeniya et al. demonstrated that treatment with a polyherbal decoction comprising *Nigella sativa* seeds, *Hemidesmus indicus* root bark, and *Smilax glabra* rhizomes significantly inhibited diethylnitrosamine (DEN)-induced hepatocarcinogenesis in male Wistar rats. The decoction markedly suppressed the development of overt hepatic tumors and prevented histopathological alterations associated with tumor formation. These findings indicate that the decoction exerts a strong chemopreventive effect against DEN-mediated liver carcinogenesis [22].

The same polyherbal decoction was further evaluated for systemic toxicity in Wistar rats and ICR mice following repeated oral administration for a period of three months. Daily administration at dose levels of 4 g/kg and 6 g/kg body weight did not result in treatment-related adverse effects on hepatic function, hematological parameters, reproductive function, or histopathological architecture of major organs, including the liver, heart, lungs, stomach, duodenum, and kidneys. In addition, no mortality or clinical signs of toxicity were observed even at an extremely high dose level of 240 g/kg/day. These findings indicate the absence of observable adverse effects over prolonged exposure and demonstrate a wide margin of safety for the decoction [23]. The safety profile observed for the optimized formulation, Vernolac, which incorporates two additional plant components, is consistent with the findings of these previously reported studies of the original polyherbal decoction.

Histopathological evaluation of the liver, spleen, kidney, lung, heart, and brain in rats from the acute toxicity study indicated that Vernolac did not cause any treatment-related organ toxicity in Wistar rats. Consequently, the LD_50_ of orally administered Vernolac exceeded 2000 mg/kg body weight in both sexes. The LD_50_ is defined as the dose of a substance that results in mortality in 50 % of animals in a group when administered orally [24]. The results of the acute toxicity study indicate that Vernolac exhibits no toxicity at a dose of 2000 mg/kg. According to OECD Guideline 420, which evaluates the acute oral toxicity of substances, if no deaths or clinical signs of toxicity are observed at a specific dose, the substance is considered to have low toxicity. Under the Global Harmonized System (GHS), if no deaths or severe clinical signs are observed at the highest dose tested (typically 2000 mg/kg), the substance is classified as Category 5 (not harmful) [25, 26].

A 28-day repeated-dose oral toxicity study was conducted in rats in accordance with OECD Guideline 407. Vernolac was administered orally at dose levels of 165, 330, and 660 mg/kg body weight per day, with distilled water used as the vehicle control. No treatment-related differences were observed in food and water consumption or relative organ weights in either sex when compared with the control groups. A statistically significant increase in body weight was observed only in male rats in the 165 mg/kg group relative to controls; however, this group had a slightly higher mean body weight at baseline at the time of randomization, and no similar effect was observed at higher dose levels. In the absence of a dose-dependent trend, the observed body weight difference was considered incidental and not related to Vernolac administration.

In female rats, hematological analysis revealed a mild but significant change in MCH at the highest dose (660 mg/kg), although values remained within the normal reference range [27]. Biochemical assessments included AST, ALT, blood urea, and serum creatinine. A significant reduction in ALT was observed only in the highest-dose Vernolac-treated female rats compared to the controls. In male rats, significant changes were noted in eosinophils, hemoglobin, HCT, MCH, RDW, and platelets, but all values remained within normal limits [27]. No significant differences were detected in the biochemical parameters of male rats compared to controls.

The liver and kidneys are the primary organs involved in the metabolism and elimination of xenobiotics [28, 29,30]. Accordingly, evaluation of hepatic and renal function is a critical component of toxicity assessment studies. In the present study, the absence of treatment-related alterations in biochemical parameters indicates that repeated oral administration of Vernolac did not produce adverse effects on liver or kidney function.

## 5. Conclusion

In the acute oral toxicity study, a single administration of Vernolac at a dose of 2000 mg/kg body weight produced no mortality or treatment-related clinical signs, indicating that the LD₅₀ is greater than 2000 mg/kg body weight via the oral route. In the 28-day repeated-dose toxicity study, daily oral administration of Vernolac at 165, 330, and 660 mg/kg did not produce toxic effects in treated Wistar rats compared with their control groups. Collectively, these findings indicate that Vernolac is well-tolerated and non-toxic at doses up to four times the therapeutic daily dose following repeated oral administration for 28 days in Wistar rats.

